# Large-domain histology-based diffusion MRI simulation via independent local simulations

**DOI:** 10.64898/2026.05.11.724295

**Authors:** Iris A. Kohler, Lei Zheng, Tristan A. Kuder, Oliver Gödicke, Mark E. Ladd, Jürgen Hesser

## Abstract

Diffusion MRI simulations based on realistic tissue microstructure provide a means to validate biophysical models and optimize acquisition protocols, but their computational cost restricts most studies to domains far smaller than a clinical voxel. The objective of this study was to develop an automated and scalable framework that converts whole-slide histology into diffusion MRI simulations at clinically relevant spatial scales while remaining feasible on standard workstation hardware. We present an end-to-end pipeline integrating two-dimensional whole-slide cell segmentation, mesh generation, and finite element Bloch-Torrey simulation. To enable simulations at large spatial scales without prohibitive memory growth, we introduce a subdomain tiling strategy in which the tissue domain is partitioned into extended subdomains simulated independently under no-flux boundary conditions. Signals are aggregated only from the central regions of each subdomain to minimize boundary artifacts. For an 800 µm × 800 µm histology-based domain, the aggregated signal differed by 0.07% from the corresponding full-domain finite element simulation while reducing wall-clock time from several days to hours and maintaining bounded memory usage independent of global domain size. When applied to a 2016 µm × 2016 µm heterogeneous region approximating the in-plane dimensions of a clinical voxel, the apparent diffusion coefficient obtained from the full domain differed from values computed in smaller dense and sparse subregions, demonstrating the influence of structural heterogeneity at clinically relevant scales on derived diffusion metrics. The proposed framework establishes an automated and memory-stable approach for generating diffusion MRI simulations directly from routine histology.

## 1. Introduction

Diffusion magnetic resonance imaging (dMRI) is a non-invasive imaging technique that probes the microscopic diffusion of water molecules in biological tissue. Quantitative interpretation of dMRI signals aims to infer biologically meaningful properties such as cellular density, extracellular volume fraction, or fiber organization [1, 2, 3, 4]. However, the spatial resolution of clinical dMRI is typically on the millimeter scale [5], far coarser than the cellular structures that govern diffusion behavior. Consequently, each imaging voxel reflects a spatially averaged signal arising from complex cellular architecture. Histology, by contrast, directly visualizes tissue microstructure at cellular resolution. Establishing a quantitative link between cellular-scale histology and millimeter-scale diffusion measurements therefore remains a central challenge in microstructural imaging.

Direct comparison between in vivo dMRI and ex vivo histology is complicated by tissue deformation, shrinkage, and staining artifacts introduced during fixation and sectioning [6], as well as by the inherent difficulty of multimodal image registration [7]. Numerical simulation offers an alternative: by generating synthetic diffusion signals directly from histology-derived tissue geometries, one can systematically investigate how specific microstructural features influence measurable diffusion metrics under controlled acquisition settings.

Several studies have demonstrated the feasibility of histology-based diffusion simulation. Two-dimensional segmentations have been extruded into simplified 3D substrates [8, 9, 10], while higher-resolution modalities such as electron microscopy [11] or confocal microscopy [12] have enabled reconstruction of detailed 3D meshes. In oncological applications, Grigoriou et al. [13] introduced a histology-informed simulation framework in which manually segmented 2D hematoxylin-eosin images were extruded to form virtual 3D environments for diffusion signal dictionary generation. These works highlight the promise of histology-driven modeling, but they often rely on manual segmentation steps, substantial computational resources, or simulations at limited spatial extents.

Even when realistic geometries are available, computational scalability poses an additional challenge. Numerical simulation methods capable of handling realistic geometries include Monte Carlo (MC) approaches, which track random walkers within digital substrates, and partial differential equation (PDE) solvers that directly solve the Bloch-Torrey equation. Representative tools include Camino [14], Disimpy [15], SpinDoctor [16], and MATI [17]. While these frameworks enable high-fidelity modeling, their computational and memory demands increase steeply with domain size and spatial resolution. As a consequence, many simulations are performed on spatial domains considerably smaller than a typical clinical voxel.

This scale discrepancy can introduce systematic bias in derived diffusion metrics. For example, [18] demonstrated that using undersized domains in randomly packed cylinder models led to biased estimates of radial signal decay. Largescale simulations approaching millimeter dimensions have been achieved using GPU acceleration or distributed computing clusters [19, 20, 21], but such approaches typically require substantial hardware infrastructure.

In this work, we address these challenges by presenting an end-to-end, scalable framework for histology-based diffusion MRI simulation. Our approach integrates two-dimensional whole-slide cell segmentation, mesh generation, and finite element Bloch-Torrey simulation into a unified automated pipeline. To enable simulations at clinically relevant in-plane voxel dimensions without prohibitive memory growth, we introduce a memory-stable subdomain tiling strategy: Large tissue regions are partitioned into non-overlapping subdomains, each of which is expanded with a fixed margin and simulated independently under no-flux boundary conditions. Diffusion signals are aggregated from the interior regions to control boundary artifacts. Because peak memory consumption depends only on subdomain size rather than total domain extent, simulations of large histological fields of view can be performed on standard workstation hardware without GPU acceleration or distributed computing.

### 1.1. Related work

#### 1.1.1. Numerical simulation frameworks and acceleration strategies

A variety of tools have been developed to simulate dMRI signals based on realistic tissue geometries. Monte Carlo (MC) methods remain the most widely used. Examples include Camino [14], which also incorporates domain optimization strategies to enable larger-scale simulations [22], the Diffusion Microscopist Simulator [23], and several GPU-accelerated implementations such as RMS [24], MC/DC [18], Disimpy [15], and SpinWalk [25]. Alternative approaches solve the Bloch-Torrey equation directly using finite element or finite difference formulations, as in SpinDoctor [16] and MATI [17]. Whereas GPU acceleration and parallelization have improved performance, none of these tools provides a general strategy for simulations on the scale of typical clinical dMRI measurements, which are constrained by both memory and computational cost. For example, our own experiments with SpinDoctor show that simulating a single 2D domain of size 1 mm × 1 mm at fine mesh resolution can exceed 1 TB of memory.

#### 1.1.2. Large-scale simulations

Several studies have extended the scale of dMRI simulations toward millimeter-sized domains using Monte Carlo tools. For instance, Jing et al. [19] modeled cardiac tissue over a 3D volume of 2500 µm × 3500 µm × 500 µm using a distributed computing cluster of 25 nodes with 8 to 32 cores and 16 to 128 GB of RAM per node, for a total of 300 cores. Rafael-Patino et al. [18] performed Monte Carlo simulations of axonal geometries spanning 1200 µm × 240 µm × 480 µm, executed over eight nodes (48 GB total RAM), and later in [20] simulated a full cubic millimeter of white matter using the GPU-accelerated MC/DC framework. Other notable large-scale examples include Yeh et al. [23] (up to 400 µm × 400 µm × 1000 µm), Bates et al. [26] (up to 500 µm voxel edge length), and Rose et al. [21], who simulated cardiac tissue over a 2.8 mm × 2.8 mm × 8 mm volume, using up to 50 compute nodes, each with two processors (10 cores per processor) and 128 GB of RAM. More recently, Wang et al. [27] modeled diffusion in neonatal cardiac tissue using parametric microstructure models within 1.38 mm × 1.38 mm × 1.4 mm voxels. The compute resources reported in these studies should not be interpreted as required for a single voxel. It is possible that authors simulated multiple voxels in parallel to explore a range of microstructural parameters.

These studies demonstrate that clinically relevant spatial scales are achievable using high-performance computing resources. However, they also illustrate the substantial computational infrastructure often required to reach such scales, including multi-node clusters or GPU acceleration. In addition, large-scale simulations are frequently performed on preconstructed or manually prepared geometries, rather than as part of an automated end-to-end workflow. Consequently, there remains a need for simulation strategies that decouple global domain size from peak memory consumption and enable clinically relevant spatial extents within accessible workstation environments.

#### 1.1.3. Boundary artifacts in diffusion MRI simulation

Diffusion MRI simulations are necessarily confined to finite spatial domains. Both these artificial boundaries as well as real diffusion barriers in biological tissue (e.g., cell membranes, myelin sheaths) give rise to the so-called “edge enhancement” described by de Swiet [28]. Thus in simulations, when employing no-flux boundary conditions to prevent spin escape at the simulation-domain boundary, the resulting magnetization field can be distorted near the domain edges. Specifically, signal attenuation is reduced relative to the interior. Several studies have mitigated this artifact by restricting signal readout to the central portion of the simulated domain, thereby excluding boundary-affected regions [29, 26]. The Monte Carlo simulator Camino [14] similarly adopts a default strategy of computing signal from only the central 75% of the substrate.

To date, however, this effect has primarily been treated as a source of bias to be minimized. To the best of our knowledge, no prior work has leveraged boundary-induced signal behavior as a means to enable more efficient or scalable large-domain diffusion MRI simulations.

#### 1.1.4. Histology-based diffusion MRI simulation

A number of prior studies have used histology-derived tissue geometries as substrates for dMRI simulation. For example, several works have relied on 2D histological segmentations of white matter, which were extruded along the third dimension to form simplified 3D substrates resembling cylindrical axons [8, 9, 10]. Others have leveraged higher-resolution microscopy: for instance, axonal geometries semi-automatically segmented from 3D electron microscopy volumes have been used directly as input for Monte Carlo simulations [11]. Confocal multiphoton microscopy has also been employed to acquire serial optical sections at approximately 1 µm thickness from plant tissue, followed by reconstruction of a full 3D mesh for diffusion simulations [12]. In cardiac applications, Rose et al. [21] segmented cardiomyocytes from 2D histology, extruded them along the *z*-axis, and introduced rotational variation to approximate local fiber architecture. More recently, Grigoriou et al. [13] proposed a histology-informed Monte Carlo framework in which manually segmented 2D hematoxylin-eosin images of liver cancer biopsies were extruded to construct virtual 3D tumor environments, enabling generation of diffusion signal dictionaries for microstructural parameter estimation.

Beyond direct segmentation, several studies have generated synthetic numerical phantoms informed by histology. Examples include packed-sphere models for lymph node tissue [30], packed-ellipsoid models for characterizing meningioma microstructure [31], myocardium-inspired substrates [26, 32], and white matter microstructure models based on realistic axonal packing statistics [33, 34].

Collectively, these works establish histology as a basis for diffusion MRI simulation. However, many existing approaches involve manual segmentation, specialized imaging modalities, or substantial computational resources. A fully automated pipeline that converts routine whole-slide histology into diffusion MRI simulations at clinically relevant spatial scales - while remaining feasible on standard workstation hardware - has not yet been established. Our work addresses this gap by integrating automated segmentation, mesh generation, and a memory-stable simulation strategy within a unified framework.

## 2. Methods

### 2.1. Histology to dMRI pipeline

In this work, we aim to establish a scalable and automated pipeline that links routine histological sections to diffusion MRI simulations at clinically relevant spatial scales.

The standard form of clinical histology consists of two-dimensional hematoxylin and eosin (H&E) stained sections. Although realistic dMRI simulations would ideally employ fully 3D tissue geometries, 3D histology - while advancing rapidly [35] - is not yet routinely acquired in clinical workflows. The proposed tiling strategy is broadly applicable to both 2D and 3D data. However, in the following sections we focus on 2D slices and corresponding cell geometries, which also offer substantially reduced computational cost.

The subsequent subsections describe our pipeline for automated cell segmentation using QuPath [36] and dMRI simulation using SpinDoctor [16]. Figure 1 provides an overview of the proposed pipeline.

**Figure 1:**
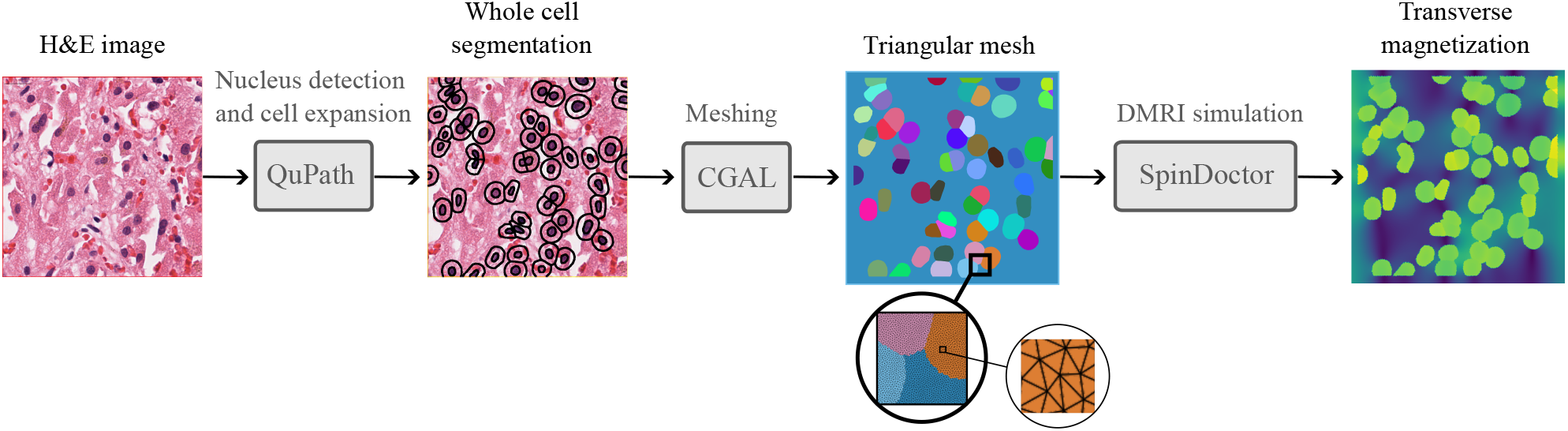
Overview of the histology-to-dMRI data pipeline. From left to right: raw H&E whole-slide image; automatic nucleus detection and cell boundary approximation performed in QuPath; triangular multicompartment mesh generated with CGAL for finite element simulation; and magnitude of the transverse magnetization field simulated with SpinDoctor.

It is important to note that each component may be replaced by analogous tools. For instance, immunofluorescence imaging may substitute for H&E staining, 3D electron microscopy may replace 2D light microscopy, and alternative dMRI simulators may be used in place of SpinDoctor.

#### 2.1.1. Whole slide images

We focus on liver tissue containing hepatocellular carcinoma (HCC). We obtained whole-slide H&E-stained liver histology slides from the publicly available TCGA-LIHC dataset [37].

#### 2.1.2. Whole cell segmentation

Our tissue model considers only intracellular space and extracellular space, excluding vasculature and subcellular detail. This simplification is consistent with prior work modeling body tissue microstructure [26, 21, 27, 38, 39].

Whole-cell segmentation was performed in QuPath [36] using a pretrained StarDist model [40]. The input image was first normalized by mapping intensities between the 1st and 99th percentiles. Cell nuclei were detected using a probability threshold of 0.4 at a resolution of 0.5 µm per pixel. Detected nuclei were then expanded radially by 5 µm to approximate whole-cell boundaries, with expansion limited to at most three times the nucleus diameter to avoid overestimating cytoplasmic extent. The final label image was resampled to a resolution of 0.1 µm/pixel to reduce discretization artifacts in rounded cell shapes.

#### 2.1.3. Mesh construction from label images

To enable finite element simulation, each 2D label image of the segmented histology patch is converted into a triangular surface mesh with per-compartment boundary markers. The resulting mesh explicitly encodes intracellular-extracellular and extracellular-external interfaces. The full mesh construction pipeline using CGAL [41] is detailed in Appendix A.

#### 2.1.4. Diffusion MRI simulation

We compute solutions of the Bloch-Torrey partial differential equation using the Julia implementation of SpinDoctor [16], which supports 2D simulations, and whhich we extended to load multi-compartment surface meshes. SpinDoctor employs the finite element method to numerically solve the Bloch-Torrey equation. The Bloch-Torrey equation models the evolution of the transverse magnetization under diffusion and relaxation. Let the simulation domain Ω be divided into *N*_*C*_ compartments (i.e. biological cells and extracellular space), denoted 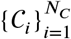. For each compartment 𝒞 _*i*_, the Bloch-Torrey equation is

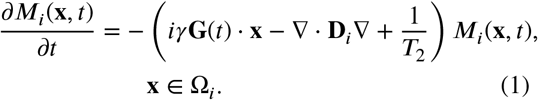

where *M*_*i*_(**x**, *t*) is the complex-valued transverse magnetization, *γ* is the gyromagnetic ratio, **G**(*t*) is the time-dependent diffusion-encoding gradient, **D**_*i*_ is the diffusion tensor, and *T*_2_ is the transverse relaxation time.

The Bloch-Torrey equation is subject to the initial conditions

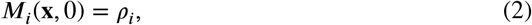

where *ρ*_*i*_ is the initial spin density of compartment 𝒞_*i*_. Further, it is subject to the interface conditions between two compartments 𝒞_*i*_ and 𝒞_*j*_

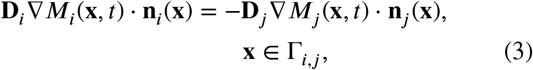

and

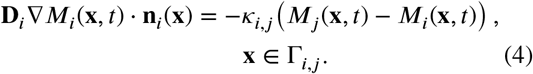

Here Γ_*i,j*_ denotes the interface between both compartments, **n**_*i*_ is the normal vector pointing outward of compartment Ω_*i*_ and *κ*_*i,j*_ is the membrane permeability coefficient.

The complex signal within a subdomain Ω_*i*_ *⊂* Ω (see Section 2.2) at echo time *T*_*E*_ is computed as

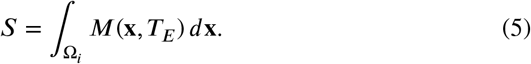

We use a standard pulsed gradient spin echo (PGSE) diffusion MRI sequence as commonly employed in clinical scanners, with parameters *b* = 800 s/mm^2^, *δ* = 9.5 ms, and Δ = 25.8 ms, corresponding to a Siemens Magnetom Vida 3 T system. Here, *δ* denotes the duration of each diffusion-encoding gradient pulse and Δ is the separation between the pulses. These parameters correspond to a diffusion-encoding gradient magnitude of approximately 74 mT/m. In the simulation, diffusion encoding is applied along the *x*-axis.

The initial spin density is set to 1, and *T*_2_ relaxation is omitted by setting *T*_2_ = *∞*. Time integration uses the Crank-Nicholson method with absolute and relative tolerances of 10^*™*6^ and 10^*™*4^, respectively.

We assume isotropic diffusion within each compartment and set the extracellular diffusivity to 0.003 mm^2^ /s, the intracellular diffusivity to 0.001 mm^2^/s and the cell membrane permeability to 0 µm/s. As experimental measurements for liver tissue were unavailable, these values were selected from ranges used in prior non-brain dMRI simulation studies [12, 26, 21, 27, 39, 38, 30, 32, 13].

SpinDoctor outputs the complex magnetization field at the mesh nodes. To compute the signal from a center crop, we interpolate the magnetization field from the triangular mesh with facet size 0.3 µm onto a regular grid with spacing 0.16 µm and numerically integrate over Ω_*i*_ using (5).

### 2.2. Scalable subdomain simulation strategy

While the above steps enable automated construction of simulation-ready tissue geometries, the resulting domains rapidly exceed feasible memory limits when scaled to clinical voxel dimensions. This section introduces a scalable simulation strategy that overcomes this limitation.

The edge-enhancement effect arising from no-flux boundary conditions at artificial simulation-domain boundaries (Section 1.1.3) is most pronounced when the characteristic diffusion length 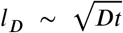, with diffusion coefficient and diffusion-encoding time *t*, remains small compared with the characteristic domain size *l*_*S*_, i.e., *l*_*D*_ ≪ *l*_*S*_ [28]. In this regime, spins originating in the interior of the domain are unlikely to encounter the boundaries during the diffusion-encoding interval and therefore do not experience the imposed no-flux constraints. Consequently, the transverse magnetization within the interior closely approximates that of an effectively unbounded medium, whereas distortions due to restricted diffusion are confined to regions near the boundaries.

We leverage this locality property to enable large-scale diffusion MRI simulations. Let Ω *⊂* ℝ^3^ denote a finite region of interest^1^ (e.g., a dMRI voxel that is too large to simulate as a single domain) that is embedded within a larger tissue domain Ω_tissue_. As illustrated in Fig. 2, the region Ω is partitioned into *N* non-overlapping subdomains 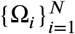. For each subdomain Ω_*i*_, an extended simulation region 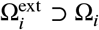 is defined by expanding Ω_i_ by a fixed margin in all spatial directions. Independent dMRI simulations are then performed on each expanded region under no-flux boundary conditions.

**Figure 2:**
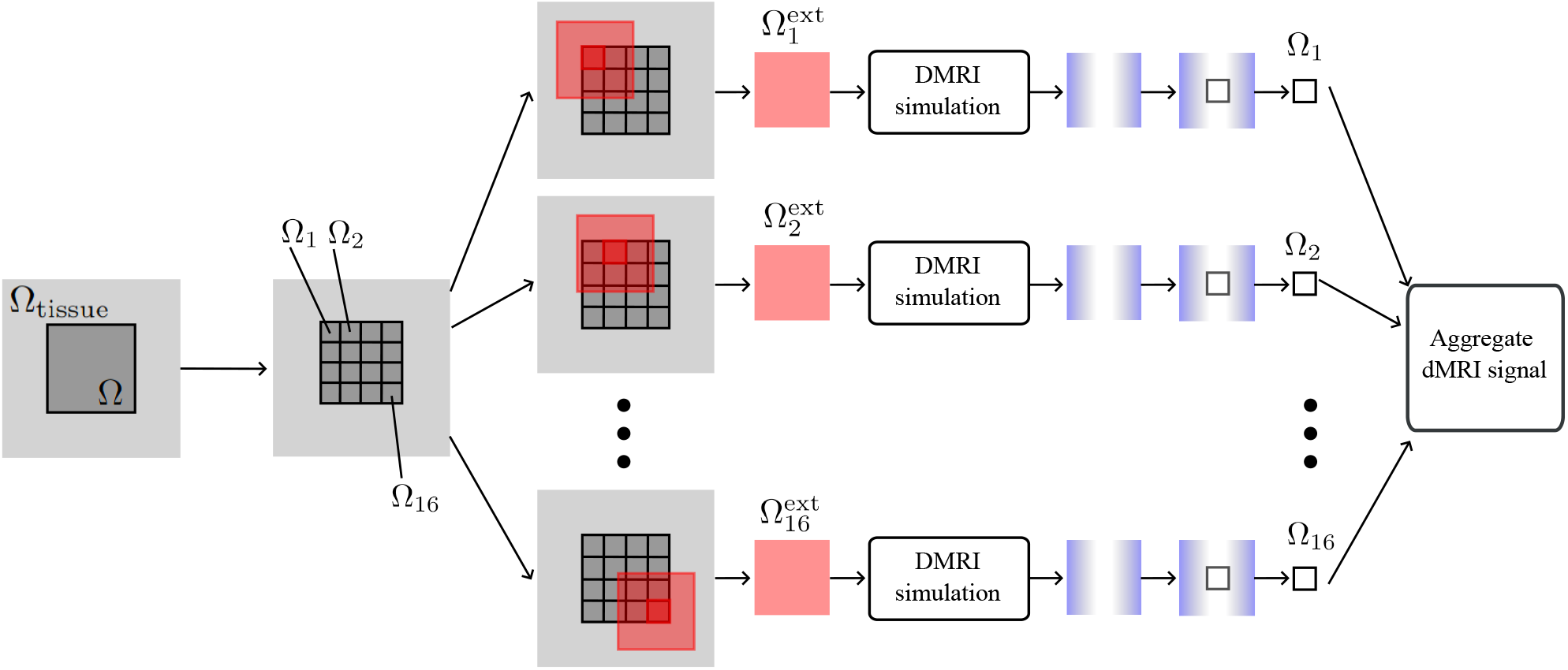
Illustration of the proposed subdomain simulation framework. Left to right: A finite region Ω is embedded in a larger domain Ω_tissue_. Ω is decomposed into non-overlapping subdomains 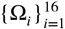. Each subdomain Ω_*i*_ is embedded in an extended simulation region 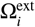. Independent simulations are performed under no-flux boundary conditions, and only the center crops corresponding to the original subdomains 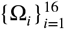 are retained. Aggregating the cropped signals approximates the diffusion-weighted signal of the original domain Ω.

Let *M*(**x**, *T*_*E*_) denote the complex transverse magnetization at spatial location **X** and echo time *T*_*E*_. After simulation, we retain only the magnetization within the central crop corresponding exactly to Ω_*i*_. The aggregated diffusion-weighted signal over the entire region Ω is then approximated as

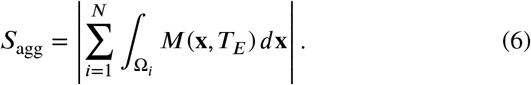

Because each subdomain Ω_*i*_ is simulated in its local coordinate system, the retained center crop represents a translated version of the same microstructure within the full domain Ω. A spatial translation by *x*_0_ introduces a global phase factor in the magnetization,

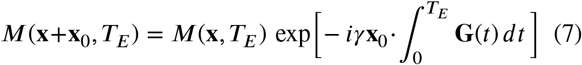

where ***G***(*t*) is the diffusion-encoding gradient and *γ* is the gyromagnetic ratio. For the diffusion MRI sequences considered in this work, namely standard pulsed-gradient spin-echo (PGSE) sequences, the gradient waveform is fully rephased, so the global phase term vanishes and the diffusion-weighted signal is invariant under translation. Consequently, the center crop of each subdomain yields the same signal as if it were embedded within the larger tissue volume.

Unlike classical overlapping domain-decomposition methods, which exchange interface data to ensure continuity across subdomains [42], our extended subdomains are simulated independently. No boundary information is shared, and the magnetization field is not enforced to be continuous. This is justified because the quantity of interest is the spatially integrated magnetization rather than the full magnetization distribution. Provided the retained center crop is sufficiently separated from the boundaries of the extended subdomain, its contribution to the aggregate signal remains effectively unaffected by the imposed boundary conditions.

Equation (6) approximates the diffusion-weighted signal in the region of interest that would arise if it were possible to simulate the entire region with boundary conditions allowing spins to interact with the surrounding tissue. The accuracy of this approximation depends on the relationship between the expansion margin and the diffusion length scale *l*_*D*_: longer diffusion times or smaller expansions increase the likelihood of boundary interactions.

### 2.3. Subdomain size and extension margin selection

To ensure that each retained crop lies sufficiently far from its simulation boundaries, we next determine suitable choices of subdomain size and expansion margin. To illustrate the procedure, and consistent with the remainder of this paper, we consider a two-dimensional example.

We empirically estimate appropriate subdomain and expansion margin dimensions by simulating diffusion in a square domain in which diffusion is unrestricted. For this idealized setting, the theoretical signal attenuation is

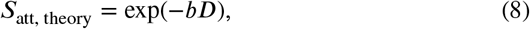

where *b* denotes the diffusion-weighting factor and *D* the diffusivity. We compute the relative error between the theoretical attenuation *S*_att, theory_ and the simulated attenuation *S*_att, sim_, evaluated over square center crops of varying sizes. The goal is to identify the largest center-crop size for which *S*_att, sim_ remains in close agreement with *S*_att, theory_.

This results in a conservative estimate of the maximum center-crop (i.e. subdomain) size. In biological tissue, membranes and organelles restrict diffusion, reducing spin displacement and diminishing boundary effects. Therefore, the selected center-crop size is expected to safely generalize to heterogeneous tissue.

Simulations were performed as described in Section 2.1.4 for square domains widths of 110 µm, 130 µm, and 150 µm. For each domain width, two simulations were performed: one with *b* = 0 and one with *b* = 800 s/mm^2^. The simulated signal attenuation was computed as

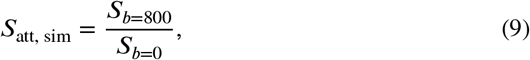

where *S*_*b*=800_ and *S*_*b*=0_ denote the absolute magnetization integrated over the cropped central region. To compute these values, the complex magnetization field was interpolated from the triangular finite element mesh (facet size 0.1 µm) to a regular grid with 0.1 µm spacing using nearest-neighbor interpolation, followed by cropping and spatial integration. The selected dimensions of the domain and the crop region resulting from this analysis are reported in Section 3.1.

### 2.4. Experimental validation and application

#### 2.4.1. Simulation of a large domain

This section presents a two-dimensional example designed to demonstrate that the proposed subdomain aggregation strategy can accurately reproduce diffusion signals obtained from a full-domain simulation, and to illustrate how the method integrates with automatic histology segmentation. Although this validation is performed in 2D for computational tractability of the reference full-domain simulation, the same framework naturally extends to 3D. However, validating the approach in 3D would require full-domain simulations at fine mesh resolution, which are prohibitively expensive with our current resources.

We compared two approaches: 1) simulating the entire domain as a single finite element mesh; 2) decomposing the domain into subdomains, simulating each extended subdomain independently with SpinDoctor, and aggregating the results by summing the signals from the center-crop regions. A histology region of interest was selected from the TCGA-LIHC whole-slide image with case ID _TCGA-UB-AA0U_, using the top-left corner coordinates (*x, y*) = (15500, 5500) µm. Cell segmentations for this region were generated in QuPath (see Section 2.1.2).

The domain size was limited by available computational resources. The largest feasible SpinDoctor simulation at high mesh resolution corresponded to a square domain of 928 µm × 928 µm, which matches a 25 × 25 grid of subdomains of size 32 µm × 32 µm, each extended to 160 µm × 160 µm. This configuration yields an effective center region of 800 µm × 800 µm that is minimally affected by boundary artifacts. Simulating domains larger than this exceeded available memory (1.1 TB RAM). For reference, typical in-plane voxel sizes in liver diffusion MRI range from 2 to 3 mm [5]. The region of interest used here is therefore smaller than a clinical voxel and serves as a constrained computational test case.

All simulations were performed with *b* = 800 s/mm^2^ using the same PGSE parameters described in Section 2.1.4. To ensure that the full-domain simulation completed within a reasonable runtime, we introduced additional parallelization into SpinDoctor by enabling parallel assembly of the finite element matrices. Specifically, we parallelized the functions `assemble_matrices, assemble_flux_matrices`, and `couple_flux_matrix` and we used 32 threads.

#### 2.4.2. Impact of domain size in heterogeneous tissue

To illustrate why simulations restricted to smaller, locally homogeneous subregions can yield misleading diffusion metrics, we constructed an example in structurally heterogeneous tissue. Diffusion parameters estimated from isolated regions may not represent the aggregate signal of a full imaging voxel. The following experiment demonstrates this effect using the proposed subdomain aggregation framework.

A 2016 µm × 2016 µm region was selected from the same TCGA-LIHC whole-slide image described in Section 2.4.1, with top-left corner coordinates (*x, y*) = (15500, 5500) µm. This dimension approximates the in-plane size of a clinical liver diffusion MRI voxel [5]. The selected region exhibited marked microstructural heterogeneity, with densely cellular areas interspersed with sparse regions.

The full domain was partitioned into square subdomains with 32 µm × 32 µm center crops and 160 µm × 160 µm extended simulation regions, consistent with Section 2.3. Independent simulations were performed on each extended region using identical mesh resolution and physical parameters. The aggregated signal of the entire 2016 µm × 2016 µm domain was computed according to (6).

Two additional 512 µm × 512 µm subregions were manually selected: a predominantly densely cellular region with top-left corner coordinates (16844, 5916) µm and a predominantly sparse region with top-left corner coordinates (16204, 6524) µm. Each subregion was independently partitioned and simulated using the same acquisition parameters and numerical settings.

All simulations were performed using a PGSE sequence with *b* = 50 s/mm^2^ and *b* = 800 s/mm^2^, with diffusion encoding applied along both the *x*-and *y*-axes, and other parameters as described in Section 2.1.4. ADC values were estimated from the logarithmic signal attenuation between the two *b*-values for each gradient direction, followed by direction averaging.

#### 2.4.3. Hardware

The reference full-domain simulations required high-memory hardware, hence all computations were performed on a dual-socket AMD EPYC 7543 system with 64 cores, 128 threads, 1.1 TB RAM, and a two-node NUMA configuration, running a 64-bit Linux operating system.

## 3. Results

### 3.1. Subdomain size and extension margin selection

To determine a subdomain size and extension margin that reliably suppress boundary artifacts while minimizing computational overhead, we simulated unrestricted diffusion in empty two-dimensional domains with widths of 110 µm, 130 µm, and 150 µm. For each domain size, we computed the simulated signal attenuation over a range of center-crop sizes and compared the results with the corresponding theoretical attenuation.

Figure 3 shows the simulated signal attenuation as a function of crop size for the three domain widths. For all domain sizes, reducing the crop size decreases the influence of boundary effects, reflected by an initial overshoot followed by a gradual flattening of the attenuation curve as the retained region moves farther from the simulation-domain boundaries. For the 110 µm domain, this plateau is not reached within the considered crop sizes, indicating that boundary effects remain present even for the smallest crop. For the 130 µm domain, an approximately flat regime emerges only at smaller crop sizes (around 20 µm), whereas for the 150 µm domain, this flattening occurs earlier and over a broader range of crop sizes.

**Figure 3:**
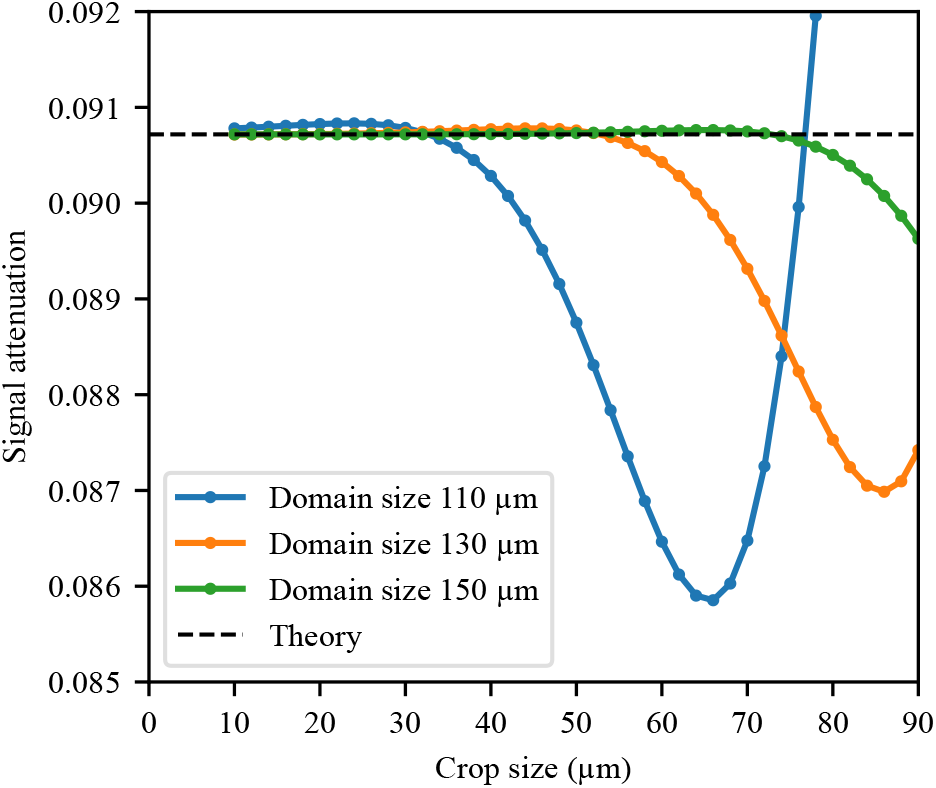
Signal attenuation as a function of crop size for unrestricted diffusion in square domains of 110 to 150 µm width, simulated at *b* = 800 s/mm^2^ with *D* = 0.003 mm^2^/s. The dashed line indicates the theoretical unrestricted diffusion signal (*S* = exp(*™bD*)).

An extended subdomain width of 150 µm combined with a center crop of 32 µm resulted in a relative error of 0.08%. This configuration^2^ was therefore selected for all subsequent experiments, as it provides a conservative margin in which convergence is achieved well within the available crop sizes.

### 3.2. Histology-based simulations

With the subdomain parameters established, we next applied the framework to histology-derived tissue geometries. Liver whole-slide histology from the TCGA-LIHC dataset was segmented to define cellular and extracellular compartments. Before performing full simulations, we first assessed mesh resolution to ensure an accurate and computationally efficient representation of the tissue microstructure.

#### 3.2.1. Mesh resolution selection

The resolution of the triangular finite-element mesh affects numerical accuracy and computational cost. We varied the resolution of the triangular mesh by adjusting the triangle size during mesh generation (see Appendix B for details).

To assess convergence behavior, which depends on cell number, size, and geometry, we conducted a mesh resolution study on 64 geometries randomly selected from the extended subdomains used to construct the large-domain simulation (Section 2.4.1). We varied the CGAL `facet_size` parameter from 1.0 to 10.0 in increments of 1.0, corresponding to approximate triangle edge lengths from 0.1 µm to 1 µm. This resulted in meshes ranging from an average of 26k to 2.23M nodes. The diffusion-weighted signals were simulated using identical settings and parameters as described in Section 2.1.4.

Simulations performed at the finest resolution (2.23M nodes) were used as a numerical reference, acknowledging that this does not constitute an analytical ground truth. For each resolution, we computed the signal errors relative to this reference.

Figure 4 summarizes the distribution of relative errors as a function of the average number of mesh nodes using boxplots, with the median relative error indicated for each resolution. Diminishing accuracy gains were observed beyond 258k nodes (corresponding to a triangle edge length of 0.3 µm). At this resolution, the median relative error was 0.05%, and all geometries exhibited errors below 0.2%.

**Figure 4:**
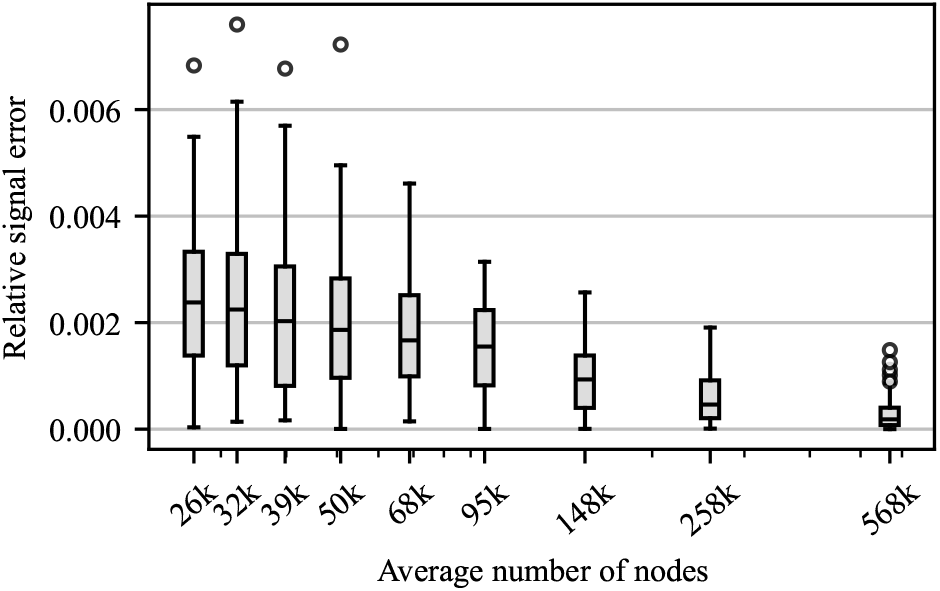
Distribution of relative dMRI signal error as a function of mesh resolution, summarized using boxplots across 64 randomly selected geometries, with the median indicated for each resolution. Errors are computed relative to the numerically most refined mesh (2.23 M nodes) and therefore reflect numerical differences rather than deviation from an analytical ground truth. The horizontal axis indicates the average number of mesh nodes for each resolution setting.

Larger errors were primarily associated with geometries containing very small cells or fine internal boundaries. Based on this accuracy-efficiency trade-off, this resolution with approximate triangle edge length of 0.3 µm was selected for all subsequent simulations.

#### 3.2.2. Simulation results of a large domain

Using the selected mesh resolution, we first evaluated the accuracy and computational cost of the proposed subdomain aggregation approach on a large region-of-interest of 928 µm × 928 µm width. Under the crop size determined in the Section 3.1, this corresponds to a 800 µm × 800 µm center-crop region unaffected by boundary effects. The region was simulated twice: first as a single full-domain simulation and second by decomposing the domain into subdomains followed by signal aggregation from the central region of the extended subdomains. Table 1 reports signal values, memory demands and runtimes for both strategies.

**Table 1.** Comparison of simulation strategies for the dMRI simulation of a 928 µm × 928 µm domain. Signal refers to the 800 µm × 800 µm center-crop region. Full-domain mesh generation was not parallelized. For the SpinDoctor simulation, the assembling of the finite element matrices was parallelized using 32 threads. For the extended subdomains, the mesh generation and simulation used 64 parallel processes. For the upper memory limit, we multiplied the maximum required memory of all individual extended subdomains (7.2 GB for mesh generation and 7.4 GB for SpinDoctor simulation) by the number of processes (64) to obtain an upper bound.

|  | Full-domain | Subdomains |
| --- | --- | --- |
| Time of mesh generation | 12 h 18 min | 4 h 48 min |
| Time of dMRI simulation | 102 h 35 min | 6 h 32 min |
| Max memory of mesh generation | 52.6 GB | 460.8 GB |
| Max memory of dMRI simulation | 869.1 GB | 473.6 GB |
| Total signal | 350 104 | 349 863 |

The full-domain simulation serves as reference. The subdomain aggregation reproduced the total signal of the 800 µm × 800 µm region with a deviation of only 0.07%, demonstrating that our approach introduces negligible numerical bias.

We note the differences in hardware usage between the two strategies. For the full-domain simulation, no parallelization was available beyond our parallel assembly of the finite element matrices (see Section 2.4.1), which utilized 32 threads to maintain a feasible runtime. In contrast, the subdomain approach enabled straightforward parallel execution across 64 processes, yielding a substantial reduction in total computation time.

Our approach also offers significant memory advantages. Although memory usage varied slightly across the 625 subdomains, reflecting differences in cell number and size, each extended subdomain required less than 7.2 GB for mesh generation and less than 7.4 GB for the SpinDoctor simulation. By comparison, the full-domain simulation demanded more than 869.1 GB of peak memory.

#### 3.2.3. Scaling of memory usage and runtime

We next evaluated how wall-clock runtime and peak memory usage scale with increasing domain size. While both mesh generation and PDE simulation contribute to the overall computational pipeline, SpinDoctor dominates the total cost. For this reason, we focus our quantitative scaling analysis on the SpinDoctor component, which represents the limiting factor for full-domain simulations.

Full-domain simulations were performed for a series of square domains, each processed using identical mesh-generation and SpinDoctor configurations. Runtime and memory usage were measured, reporting wall-clock time and peak resident memory.

Table 2 summarizes the results for domain widths ranging from 192 µm to 928 µm. Both runtime and peak memory increased sharply with domain size, reflecting the rapid growth of finite element matrix dimensions in full-domain simulations.

**Table 2.** Wall-clock runtime and peak memory usage for full-domain SpinDoctor simulations. Each simulation used 32 threads for the assembly of the finite element matrices but was otherwise single-threaded.

| Domain size | Wall-clock time | Peak memory |
| --- | --- | --- |
| $192 \mu\text{m} \times 192 \mu\text{m}$ | 41 min 50 s | 19.5 GB |
| $288 \mu\text{m} \times 288 \mu\text{m}$ | 1 h 50 min 39 s | 68.5 GB |
| $448 \mu\text{m} \times 448 \mu\text{m}$ | 5 h 21 min 54 s | 123.3 GB |
| $608 \mu\text{m} \times 608 \mu\text{m}$ | 13 h 50 min 12 s | 140.4 GB |
| $768 \mu\text{m} \times 768 \mu\text{m}$ | 35 h 38 min 02 s | 409.6 GB |
| $928 \mu\text{m} \times 928 \mu\text{m}$ | 93 h 26 min 24 s | 869.1 GB |

In contrast, our subdomain approach does not exhibit global memory growth. Each extended subdomain is simulated independently, and memory usage is effectively bounded by that of a single subdomain solve.

To visualize these trends, Fig. 5 plots domain width versus peak memory usage. For the extended subdomains, an upper bound on memory consumption is computed by multiplying the maximum observed peak memory per subdomain (i.e. 7.4 GB) by the number of parallel processes used (i.e. 64). This bound remains constant across domain sizes, whereas the full-domain memory demand increases rapidly.

**Figure 5:**
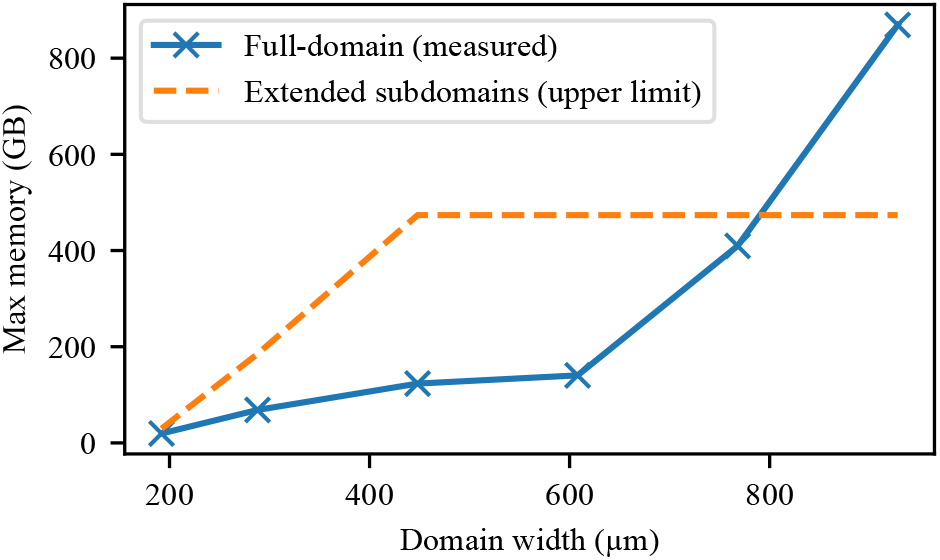
Peak memory usage as a function of domain width for full-domain SpinDoctor simulations using a single process (solid line) and an upper bound for the extended subdomains using 64 parallel processes (dashed line). Memory demand for the subdomain simulation remains effectively constant, whereas full-domain memory usage increases steeply with domain size.

The subdomain simulations exhibit embarrassingly parallel behavior and scale effectively with the number of available processing units. On our dual-socket AMD EPYC 7543 system (128 hardware threads), we achieved a sustained throughput of 96 subdomains per hour when running 64 concurrent processes. Increasing the degree of parallelism beyond this level showed diminishing returns, primarily due to memory-bandwidth constraints.

To assess the effect of memory constraints on runtime, we consider the serial scenario where subdomains are simulated one at a time. In this case, the total wall-clock time increases to approximately 444 hours, about 4.3 times longer than the full-domain simulation. With 64 parallel processes, the wall-clock time reduces to 6.5 hours, representing a 14-fold speedup relative to the full-domain simulation. These results highlight the trade-off between memory usage and computational time: subdomain decomposition substantially lowers memory requirements while enabling efficient parallel execution, though serial execution incurs a runtime penalty.

#### 3.2.4. Impact of domain size in heterogeneous tissue

Finally, to illustrate the practical impact of simulations at clinical voxel dimensions, we applied the proposed framework to a 2016 µm × 2016 µm heterogeneous region. Figure 6 shows the histology-derived domain alongside the two 512 µm × 512 µm subregions representing predominantly dense and sparse cellular environments. The full domain exhibits substantial spatial variation in cell density.

**Figure 6:**
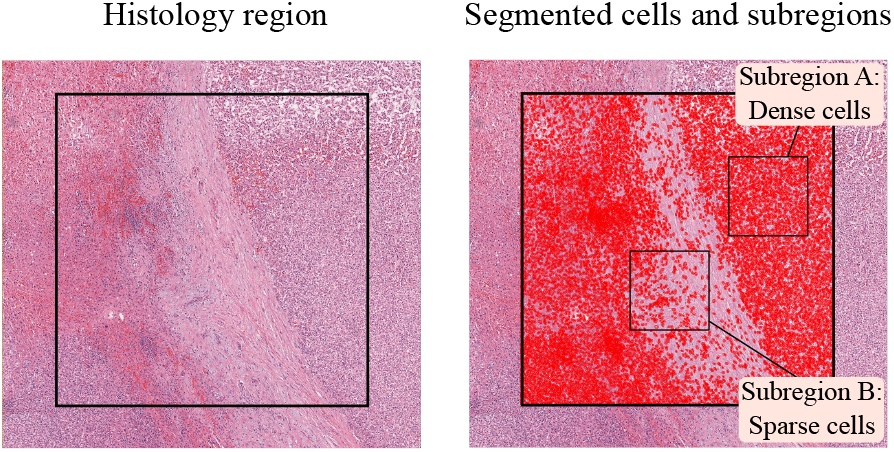
Histology domain and corresponding segmentation overlay. Left: 2016 µm × 2016 µm region extracted from the TCGA-LIHC whole-slide H&E image with case ID `TCGA-UB-AA0U`. Right: Same region with detected nuclei and corresponding whole-cell boundaries overlaid in red. Black squares indicate the 512 µm × 512 µm dense and sparse subregions used for comparison.

The direction-averaged ADC obtained from the full heterogeneous domain was 8.06 × 10^*™*4^ mm^2^/s. For the predominantly dense subregion, the ADC was 6.08 × 10^*™*4^ mm^2^/s, whereas the predominantly sparse subregion yielded an ADC of 15.09 × 10^*™*4^ mm^2^/s.

Relative to the full-domain simulation, the ADC of the dense subregion was 24.6% lower, whereas the ADC of the sparse subregion was 87.2% higher.

These findings demonstrate that small-domain simulations may misrepresent diffusion behavior at clinically relevant voxel sizes in heterogeneous tissue. The proposed subdomain aggregation framework enables simulation at spatial scales comparable to clinical voxels, thereby supporting more representative histology-based diffusion modeling.

## 4. Discussion

In this work, we introduced a scalable and automated framework that links routine histological tissue sections to diffusion MRI simulations at clinically relevant spatial scales. While histology directly visualizes cellular architecture, translating such microstructural information into diffusion signals measured over millimeter-scale imaging voxels has been computationally prohibitive due to the steep memory growth of full-domain finite element simulations. By partitioning the simulation domain into independent subdomains, extending them by a fixed margin, and aggregating the diffusion-weighted signal only from center-cropped regions, the proposed method enables large-scale simulations that closely approximate an unbounded medium while maintaining bounded memory usage. Applied to a histology-derived liver dataset, the framework reproduced the diffusion signal of an 800 µm × 800 µm region with negligible deviation from a full-domain reference. These results demonstrate that aggregation of local simulations preserves numerical fidelity in heterogeneous, biologically realistic microstructure and provides a practical forward model from cellular architecture to millimeter-scale dMRI signals.

Beyond demonstrating numerical fidelity, the large-domain experiment indicates that scalability has implications beyond computational efficiency. In structurally heterogeneous tissue, the diffusion-weighted signal of a voxel represents a composite response arising from multiple microstructural environments and cannot, in general, be inferred from simulations of small homogeneous subregions. In our 2016 µm × 2016 µm example, the ADC differed measurably from the ADC obtained from either the predominantly dense or predominantly sparse regions alone. This observation suggests that limited-domain simulations may introduce bias when diffusion metrics are used for microstructure model validation or for establishing histology-MRI correspondences. By enabling simulations at spatial scales comparable to clinical voxels while preserving underlying heterogeneity, the proposed framework supports more representative digital phantom construction and more robust evaluation of diffusion modeling approaches.

From a computational perspective, an important advantage of the proposed formulation is its natural amenability to parallelization. Whereas full-domain simulations are limited by mesh size, global matrix assembly, and memory overhead, the subdomain simulations can be distributed across multiple processes with minimal communication cost. In our experiments, running the 625 extended subdomains on 64 parallel processes reduced total runtime from several days to hours and eliminated peak memory usage concerns. This suggests that our approach may provide a practical route toward routine high-resolution dMRI simulations on large tissue sections without reliance on specialized high-memory compute nodes.

Although the present study focused on two-dimensional histological sections, the underlying concept is directly extensible to 3D simulations. The edge-enhancement effect that motivates the center-cropping arises from fundamental properties of restricted diffusion and therefore also applies in 3D. Extending the framework to volumetric histology, 3D microscopy, or reconstructed tissue stacks could provide more efficient simulations of realistic digital phantoms for validating biophysical models.

The approach, however, has limitations. Its validity depends on diffusion times being short relative to the characteristic domain scale. At long diffusion times, spins explore a larger region of space and are more likely to interact with subdomain boundaries, requiring substantially larger subdomains to maintain an artifact-free central region. Whereas such long diffusion times are uncommon in clinical dMRI due to reduced signal-to-noise ratio, they may arise in preclinical or research settings, in which case the computational savings of the method would diminish.

A further practical limitation concerns scalability with respect to the number of parallel processes. Although the subdomain simulations are theoretically embarrassingly parallel, we observed diminishing returns when increasing the number of concurrent processes beyond 64 on our test system. This behavior was attributable to memory-bandwidth limitations. In computing environments with higher bandwidth or distributed-memory architectures (e.g., multi-node clusters), such limitations may be substantially reduced or absent. Thus, while the method scales favorably in principle, practical performance will depend on the characteristics of the underlying hardware.

Future work may further expand the utility of the approach. Although the large-domain experiments in this study were performed on a high-memory multi-socket system to enable direct comparison with full-domain simulations, the proposed subdomain formulation itself does not require such hardware. Each extended subdomain can be simulated independently with a memory footprint below 8 GB, making the framework executable on conventional CPU-based workstations, albeit with longer wall-clock times in serial execution. Integration with GPU-accelerated Monte Carlo tools such as MC/DC [18] could further accelerate simulations for large parameter sweeps or high-throughput studies. Importantly, the subdomain formulation is agnostic to the underlying numerical solver and can be combined with either finite element or Monte Carlo implementations. Improvements in automated histology segmentation, such as deep-learning-based 3D cell boundary detection or reconstruction from multi-slice stacks, would enhance the biological realism of input geometries. Together, these directions could enable high-throughput generation of histology-based diffusion phantoms, facilitating systematic evaluation of microstructure models, acquisition design, and interpretation of diffusion biomarkers across a wide range of tissues and disease contexts.

## 5. Conclusion

We presented an automated and scalable framework for histology-based diffusion MRI simulation that enables simulations at spatial scales approaching those of clinical imaging voxels. By combining whole-slide cell segmentation, mesh generation, finite element Bloch-Torrey simulation, and a subdomain aggregation strategy, the proposed approach overcomes the prohibitive memory growth associated with conventional full-domain simulations.

The results demonstrate that aggregating signals from independently simulated extended subdomains reproduces full-domain diffusion signals with high accuracy while maintaining bounded memory usage independent of global domain size. Applied to heterogeneous liver histology, the framework further showed that diffusion metrics derived from small locally homogeneous regions can differ substantially from those obtained at clinically relevant spatial scales, underscoring the importance of preserving tissue heterogeneity in histology-based diffusion modeling.

Beyond the specific experiments presented here, the proposed formulation is general and compatible with different segmentation strategies and numerical solvers. Although this study focused on two-dimensional histology, the same principles naturally extend to volumetric tissue reconstructions and three-dimensional diffusion simulations. The framework therefore provides a practical foundation for large-scale histology-informed digital phantom generation, quantitative validation of biophysical diffusion models, and systematic investigation of the relationship between tissue microstructure and diffusion MRI measurements.

## A. Appendix: Mesh generation

SpinDoctor requires multi-compartment 2D triangular meshes as input. Existing tools for meshing from 2D label images often fail when three or more compartments meet at a point or share edges. CGAL [41] can robustly resolve such configurations, but only in 3D. We therefore generate a 3D mesh by extrusion and subsequently extract the required 2D mesh.

Prior to meshing, background regions smaller than five pixels are removed to prevent disconnected fragments. Each image is padded by 5 µm on all sides and assigned to the extracellular space (ECS) to avoid complications with the mesh generation and simulation.

To apply CGAL’s `make_mesh_3` function, the 2D image is stacked along the *z*-axis and converted to INR format. Mesh resolution is primarily controlled by the `facet_size`parameter, with the facet angle fixed at 30° and the cell radius-edge ratio set to 1.5 to ensure well-shaped tetrahedra. Lloyd smoothing is applied post-meshing to improve surface quality.

A 2D surface mesh is extracted by region growing from a seed triangle on the *z* = 0 plane. Coplanar triangles are added unless they lie within one voxel of the top or bottom cuboid boundaries, which avoids selecting side faces. Boundary markers are assigned such that the outer ECS boundary and each unique inter-compartment interface receive distinct labels. Disconnected interfaces between the same compartment pair share a marker, consistent with SpinDoctor conventions.

Disconnected ECS components are identified and assigned unique region labels. Isolated triangle fragments, typically attached by a single node due to meshing artifacts, are relabeled as ECS. Finally, the mesh is rescaled so that one unit corresponds to 1 µm, enabling use of label images at arbitrary resolutions after appropriate scaling.

## B. Appendix: Mesh resolution parameters

Mesh resolution trades off geometric accuracy and computational cost. We varied resolution through CGAL’s `facet_size` parameter. Because other meshing constraints may implicitly increase triangle size, related parameters were adjusted to ensure that effective surface edge lengths did not exceed the target `facet_size.` Table 3 summarizes the parameter combinations used to generate progressively finer meshes.

**Table 3.** Parameter combinations for mesh generation. Facet size bounds surface triangle edge length; cell size limits tetrahedral cell size; facet distance controls the maximum deviation between the true and meshed surfaces. The facet angle (minimum surface triangle angle) was fixed at 30^°^, and the cell radius-edge ratio (upper bound on circumsphere radius to shortest edge) was fixed at 1.5 for all meshes. Stack count denotes the number of image stacks used for extrusion along the *z*-axis, and tolerance specifies the distance threshold for selecting triangles near the *z* = 0 plane when extracting the 2D surface.

| Facet Size | Cell Size | Facet Distance | Stack Count | Tolerance |
| --- | --- | --- | --- | --- |
| 10.0 | 10.0 | 5.0 | 50 | 5.0 |
| 9.0 | 9.0 | 4.5 | 50 | 5.0 |
| 8.0 | 8.0 | 4.0 | 30 | 4.0 |
| 7.0 | 7.0 | 3.5 | 30 | 4.0 |
| 6.0 | 6.0 | 3.0 | 30 | 3.5 |
| 5.0 | 5.0 | 2.5 | 20 | 2.0 |
| 4.0 | 4.0 | 2.0 | 20 | 2.0 |
| 3.0 | 3.0 | 1.5 | 20 | 1.5 |
| 2.0 | 2.0 | 1.0 | 10 | 1.5 |
| 1.0 | 1.0 | 0.5 | 10 | 1.0 |

## CRediT authorship contribution statement

**Iris A. Kohler:** Conceptualization, Data curation, Formal analysis, Investigation, Methodology, Software, Validation, Visualization, Writing - original draft, Writing - review & editing. **Lei Zheng:** Project administration, Visualization, Writing - review & editing. **Tristan A. Kuder:** Writing - review & editing, Funding acquisition. **Oliver Gödicke:** Writing - review & editing. **Mark E. Ladd:** Funding acquisition, Writing - review & editing. **Jürgen Hesser:** Conceptualization, Funding acquisition, Project administration, Supervision, Writing - review & editing.

## Data availability

The code for mesh generation is available at https://github.com/irisakohler/histology-image-to-mesh. The modified SpinDoctor code is at https://github.com/irisakohler/SpinDoctor.jl-custom-mesh.

## Acknowledgment

This research was supported by the CZS Heidelberg Initiative for Model-Based AI (MBAI) under the Grant P2021-02-001. The authors gratefully acknowledge their support. The authors gratefully acknowledge the data storage service SDS@hd supported by the Ministry of Science, Research and the Arts Baden-Württemberg (MWK) and the German Research Foundation (DFG) through grant INST 35/1503-1 FUGG.

The results shown here are in whole or part based upon data generated by the TCGA Research Network: https://www.cancer.gov/tcga.

## Footnotes

1 For clarity, we describe the general 3D formulation. All experiments in this work are performed in 2D.

2 Note that the mesh generation process adds 5 µm of extracellular padding on all sides, resulting in an actual size of the extended subdomain of 160 µm × 160 µm.

## References

[1] E. Fokkinga, J. A. Hernandez-Tamames, A. Ianus, M. Nilsson, C. M. Tax, R. Perez-Lopez, F. Grussu, Advanced diffusion-weighted MRI for cancer microstructure assessment in body imaging, and its relationship with histology, Journal of Magnetic Resonance Imaging 60 (2023) 1278–1304.

[2] E. Martinez-Heras, F. Grussu, F. Prados, E. Solana, S. Llufriu, Diffusion-weighted imaging: Recent advances and applications, Seminars in Ultrasound, CT and MRI 42 (2021) 490–506.

[3] M. Afzali, T. Pieciak, S. Newman, E. Garyfallidis, E. Özarslan, H. Cheng, D. K. Jones, The sensitivity of diffusion MRI to microstructural properties and experimental factors, Journal of Neuroscience Methods 347 (2021). Art. no. 108951.

[4] I. O. Jelescu, M. Palombo, F. Bagnato, K. G. Schilling, Challenges for biophysical modeling of microstructure, Journal of Neuroscience Methods 344 (2020). Art. no. 108861.

[5] A. Shukla-Dave, N. A. Obuchowski, T. L. Chenevert, S. Jambawalikar, L. H. Schwartz, D. Malyarenko, W. Huang, S. M. Noworolski, R. J. Young, M. S. Shiroishi, H. Kim, C. Coolens, H. Laue, C. Chung, M. Rosen, M. Boss, E. F. Jackson, Quantitative imaging biomarkers alliance (QIBA) recommendations for improved precision of DWI and DCE-MRI derived biomarkers in multicenter oncology trials, Journal of Magnetic Resonance Imaging 49 (2019) e101–e121.

[6] L. Alic, J. C. Haeck, S. Klein, K. Bol, S. T. van Tiel, P. A. Wielopolski, M. Bijster, W. J. Niessen, M. Bernsen, J. F. Veenland, M. de Jong, Multi-modal image registration: matching MRI with histology, in: Medical Imaging 2010: Biomedical Applications in Molecular, Structural, and Functional Imaging, volume 7626, SPIE, 2010. doi:10.1117/12.844123, Art. no. 762603.

[7] A. Sen, P. Troncoso, A. Venkatesan, M. D. Pagel, J. A. Nijkamp, Y. He, A. C. Lesage, M. Woodland, K. K. Brock, Correlation of in-vivo imaging with histopathology: A review, European Journal of Radiology 144 (2021). Art. no. 109964.

[8] C. L. Chin, F. W. Wehrli, S. N. Hwang, M. Takahashi, D. B. Hackney, Biexponential diffusion attenuation in the rat spinal cord: Computer simulations based on anatomic images of axonal architecture, Magnetic Resonance in Medicine 47 (2002) 455–460.

[9] C. L. Chin, F. W. Wehrli, Y. Fan, S. N. Hwang, E. D. Schwartz, J. Nissanov, D. B. Hackney, Assessment of axonal fiber tract architecture in excised rat spinal cord by localized NMR q-space imaging: Simulations and experimental studies, Magnetic Resonance in Medicine 52 (2004) 733–740.

[10] J. Xu, H. Li, K. D. Harkins, X. Jiang, J. Xie, H. Kang, M. D. Does, J. C. Gore, Mapping mean axon diameter and axonal volume fraction by MRI using temporal diffusion spectroscopy, NeuroImage 103 (2014) 10–19.

[11] H. H. Lee, K. Yaros, J. Veraart, J. L. Pathan, F. X. Liang, S. G. Kim, D. S. Novikov, E. Fieremans, Along-axon diameter variation and axonal orientation dispersion revealed with 3D electron microscopy: implications for quantifying brain white matter microstructure with histology and diffusion MRI, Brain Structure and Function 224 (2019) 1469–1488.

[12] E. Panagiotaki, M. G. Hall, H. Zhang, B. Siow, M. F. Lythgoe, D. C. Alexander, High-fidelity meshes from tissue samples for diffusion MRI simulations, in: Medical Image Computing and Computer-Assisted Intervention – MICCAI 2010, Springer Berlin Heidelberg, Berlin, Heidelberg, 2010, pp. 404–411.

[13] A. Grigoriou, C. Macarro, M. Palombo, D. Navarro-Garcia, A. K. Voronova, K. Bernatowicz, I. Barba, A. Escriche, E. Greco, M. Abad, S. Simonetti, G. Serna, R. Mast, X. Merino, N. Roson, M. Escobar, M. Vieito, P. Nuciforo, R. Toledo, E. Garralda, R. Sala-Llonch, E. Fieremans, D. S. Novikov, R. Perez-Lopez, F. Grussu, Histology-informed microstructural diffusion simulations for MRI cancer characterisation—the Histo-µSim framework, Communications Biology 8 (2025). Art. no. 1695.

[14] P. A. Cook, Y. Bai, S. Nedjati-Gilani, K. K. Seunarine, M. G. Hall, G. J. Parker, D. C. Alexander, Camino: Open-source diffusion-MRI reconstruction and processing, in: Proceedings of the 14th Scientific Meeting of the International Society for Magnetic Resonance in Medicine (ISMRM), Seattle, WA, USA, 2006, p. 2759.

[15] L. Kerkelä, F. Nery, M. Hall, C. Clark, Disimpy: A massively parallel Monte Carlo simulator for generating diffusion-weighted mri data in python, Journal of Open Source Software 5 (2020). Art. no. 2527.

[16] J. R. Li, V. D. Nguyen, T. N. Tran, J. Valdman, C. B. Trang, K. V. Nguyen, D. T. S. Vu, H. A. Tran, H. T. A. Tran, T. M. P. Nguyen, SpinDoctor: A MATLAB toolbox for diffusion MRI simulation, NeuroImage 202 (2019). Art. no. 16120.

[17] J. Xu, S. P. Devan, D. Shi, A. Pamulaparthi, N. Yan, Z. Zu, D. S. Smith, K. D. Harkins, J. C. Gore, X. Jiang, MATI: A GPU-accelerated toolbox for microstructural diffusion MRI simulation and data fitting with a graphical user interface, Magnetic Resonance Imaging 122 (2025). Art. no. 110428.

[18] J. Rafael-Patino, D. Romascano, A. Ramirez-Manzanares, E. J. Canales-Rodríguez, G. Girard, J. P. Thiran, Robust Monte-Carlo simulations in Diffusion-MRI: Effect of the substrate complexity and parameter choice on the reproducibility of results, Frontiers in Neuroinformatics 14 (2020). Art. no. 8.

[19] Y. Jing, I. E. Magnin, C. Frindel, Monte Carlo simulation of water diffusion through cardiac tissue models, Medical Engineering and Physics 120 (2023). Art. no. 104013.

[20] J. Rafael-Patino, G. Girard, R. Truffet, M. Pizzolato, E. Caruyer, J.-P. Thiran, The diffusion-simulated connectivity (DiSCo) dataset, Data in Brief 38 (2021). Art. no. 107429.

[21] J. N. Rose, S. Nielles-Vallespin, P. F. Ferreira, D. N. Firmin, A. D. Scott, D. J. Doorly, Novel insights into in-vivo diffusion tensor cardiovascular magnetic resonance using computational modeling and a histology-based virtual microstructure, Magnetic Resonance in Medicine 81 (2019) 2759–2773.

[22] M. G. Hall, G. Nedjati-Gilani, D. C. Alexander, Realistic voxel sizes and reduced signal variation in Monte Carlo simulation for diffusion MR data synthesis, arXiv:1701.03634, 2017.

[23] C. H. Yeh, B. Schmitt, D. L. Bihan, J. R. Li-Schlittgen, C. P. Lin, C. Poupon, Diffusion Microscopist Simulator: A general Monte Carlo simulation system for diffusion magnetic resonance imaging, PLoS ONE 8 (2013). Art. no. e76626.

[24] H. H. Lee, E. Fieremans, D. S. Novikov, Realistic Microstructure Simulator (RMS): Monte Carlo simulations of diffusion in three-dimensional cell segmentations of microscopy images, Journal of Neuroscience Methods 350 (2021). Art. no. 109018.

[25] A. Aghaeifar, S. Mueller, K. Scheffler, SpinWalk: A Monte Carlo simulator for MR-signal formation in inhomogeneous tissue, Imaging Neuroscience 3 (2025). Art. no. imag_a_00533.

[26] J. Bates, I. Teh, D. McClymont, P. Kohl, J. E. Schneider, V. Grau, Monte Carlo simulations of diffusion weighted MRI in myocardium: Validation and sensitivity analysis, IEEE Transactions on Medical Imaging 36 (2017) 1316–1325.

[27] L. Wang, Y. Hong, Y. B. Qin, X. Y. Cheng, F. Yang, J. Yang, Y. M. Zhu, Connecting macroscopic diffusion metrics of cardiac diffusion tensor imaging and microscopic myocardial structures based on simulation, Medical Image Analysis 77 (2022). Art. no. 102325.

[28] T. M. de Swiet, Diffusive edge enhancement in imaging, Journal of Magnetic Resonance, Series B 109 (1995) 12–18.

[29] S. N. Hwang, C. L. Chin, F. W. Wehrli, D. B. Hackney, An image-based finite difference model for simulating restricted diffusion, Magnetic Resonance in Medicine 50 (2003) 373–382.

[30] R. Gardier, A. Savoy, J. L. V. Haro, G. Girard, E. J. Canales-Rodriguez, E. Fischi-Gomez, A. Hertanu, I. O. Jelescu, J. Rafael-Patino, J. P. Thiran, Comparing Steam and PGSE diffusion MRI signal of rat lymph nodes using in-silico simulations, in: 2024 IEEE International Symposium on Biomedical Imaging (ISBI), IEEE Computer Society, 2024, pp. 1–5. doi:10.1109/ISBI56570.2024.10635470.

[31] G. Buizza, C. Paganelli, F. Ballati, S. Sacco, L. Preda, A. Iannalfi, D. C. Alexander, G. Baroni, M. Palombo, Improving the characterization of meningioma microstructure in proton therapy from conventional apparent diffusion coefficient measurements using Monte Carlo simulations of diffusion MRI, Medical Physics 48 (2021) 1250–1261.

[32] M. Lashgari, N. Ravikumar, I. Teh, J. R. Li, D. L. Buckley, J. E. Schneider, A. F. Frangi, Three-dimensional micro-structurally informed in silico myocardium—towards virtual imaging trials in cardiac diffusion weighted MRI, Medical Image Analysis 82 (2022). Art. no. 102592.

[33] J. L. Villarreal-Haro, R. Gardier, E. J. Canales-Rodríguez, E. Fischi-Gomez, G. Girard, J. P. Thiran, J. Rafael-Patiño, CACTUS: a computational framework for generating realistic white matter microstructure substrates, Frontiers in Neuroinformatics 17 (2023). Art. no. 1208073.

[34] K. Ginsburger, F. Matuschke, F. Poupon, J.-F. Mangin, M. Axer, C. Poupon, MEDUSA: A GPU-based tool to create realistic phantoms of the brain microstructure using tiny spheres, NeuroImage 193 (2019) 10–24.

[35] X. Xu, J. Su, R. Zhu, K. Li, X. Zhao, J. Fan, F. Mao, From morphology to single-cell molecules: high-resolution 3D histology in biomedicine, Molecular Cancer 24 (2025). Art. no. 63.

[36] P. Bankhead, M. B. Loughrey, J. A. Fernández, Y. Dombrowski, D. G. McArt, P. D. Dunne, S. McQuaid, R. T. Gray, L. J. Murray, H. G. Coleman, J. A. James, M. Salto-Tellez, P. W. Hamilton, QuPath: Open source software for digital pathology image analysis, Scientific Reports 7 (2017). Art. no. 16878.

[37] B. J. Erickson, S. Kirk, Y. Lee, O. Bathe, M. Kearns, C. Gerdes, K. Rieger-Christ, J. Lemmerman, The Cancer Genome Atlas Liver Hepatocellular Carcinoma Collection (TCGA-LIHC) (Version 5) [Data set], https://doi.org/10.7937/K9/TCIA.2016.IMMQW8UQ, 2016. doi:10.7937/K9/TCIA.2016.IMMQW8UQ, The Cancer Imaging Archive.

[38] R. Gardier, J. L. V. Haro, E. J. Canales-Rodríguez, I. O. Jelescu, G. Girard, J. Rafael-Patiño, J. P. Thiran, Cellular Exchange Imaging (CEXI): Evaluation of a diffusion model including water exchange in cells using numerical phantoms of permeable spheres, Magnetic Resonance in Medicine 90 (2023) 1625–1640.

[39] F. Grussu, K. Bernatowicz, I. Casanova-Salas, N. Castro, P. Nuciforo, J. Mateo, I. Barba, R. Perez-Lopez, Diffusion MRI signal cumulants and hepatocyte microstructure at fixed diffusion time: Insights from simulations, 9.4T imaging, and histology, Magnetic Resonance in Medicine 88 (2022) 365–379.

[40] U. Schmidt, M. Weigert, C. Broaddus, G. Myers, Cell detection with star-convex polygons, in: Medical Image Computing and Computer Assisted Intervention – MICCAI 2018, volume 11071 LNCS, Springer Verlag, 2018, pp. 265–273. doi:10.1007/978-3-030-00934-2_30.

[41] The CGAL Project, CGAL User and Reference Manual, 6.0.1 ed., CGAL Editorial Board, 2024. URL: https://doc.cgal.org/6.0.1/Manual/packages.html.

[42] T. F. Chan, T. P. Mathew, Domain decomposition algorithms, Acta Numerica 3 (1994) 61–143.

